# Characterization of Gα_s_ and Gα_olf_ activation by catechol and non-catechol dopamine D1 receptor agonists

**DOI:** 10.1101/2023.10.03.560682

**Authors:** Anh Minh Nguyen, Ana Semeano, Vianna Quach, Asuka Inoue, David E. Nichols, Hideaki Yano

**Author notes:** Corresponding author: Hideaki Yano.

## Abstract

The dopamine D1 receptor (D1R) couples to Gα_s_ and Gα_olf_ and plays a crucial role in regulating voluntary movement and other cognitive functions, making it a potential therapeutic target for several neurological and neuropsychiatric disorders, such as Parkinson’s disease and schizophrenia. In the central nervous system, Gα_s_ is widely expressed in the cortex and Gα_olf_ is predominantly found in the striatum. We used two different configurations of bioluminescence resonance energy transfer (BRET) assays and a fluorescence-based cyclic AMP (cAMP) production functional assay to test a series of tetracyclic catechol (dihydrexidine, methyl-dihydrexidine, doxanthrine) and non-catechol (tavapadon, PF-8294, PF-6142) D1R agonists for their activity at these G proteins. We discovered that these tetracyclic catechol compounds, PF-8294 and PF-6142 exerted full agonism when D1R coupled to Gα_s_ but partial agonism when D1R coupled to Gα_olf_. In contrast, tavapadon acted as a full agonist at Gα_olf_ and a partial agonist at Gα_s_. The effects of these compounds on the cortical and nigral electrophysiological events agree with their selectivity profiles. This suggests the possibility of achieving region-specific pharmacology and opens new directions for developing D1R drugs to treat relevant neurological and neuropsychiatric disorders.

## INTRODUCTION

The dopamine D1 receptor (D1R) is one of the most widely expressed dopamine (DA) receptors in the mammalian central nervous system (CNS).^1^ These receptors mediate the output of most dopaminergic pathways to regulate a breadth of important physiological and behavioral functions.^2, 3^ D1R activation has been reported to stimulate locomotor activity,^4, 5^ drive motivation and reward-associated behaviors,^6, 7^ and mediate executive functions and working memory.^8–10^ Thus, D1R appears to be a promising target for drug development against several neuropsychiatric disorders, including Parkinson’s Disease (PD) and Schizophrenia.

Developing selective D1R agonists has proved to be challenging. Classic selective D1R agonists usually resemble the catechol moiety of DA. Although some D1R agonists exhibited promising effects on cognitive function^11^ and locomotor activity,^12^ with some even advanced to clinical trials,^13, 14^ they were prematurely terminated due to their severe side effects, unfavorable pharmacokinetic properties, and tolerance.^14–17^ Because of these issues, there is a dire need for new D1R agonists that can overcome these limitations but are equally efficacious. In recent years, a new class of D1R agonists without the catechol moiety has been discovered.^18–21^ Remarkably, a few ligands from this class have demonstrated antiparkinsonian effects, encouraging pharmacokinetic profiles, and limited tolerance.^22–27^ However, little is known about the pharmacological differences since the conventional catechol and novel non-catechol agonists have not been compared in various assay modalities in a systematic way.

Among the transducers that couple to D1R, two related Gα_s_-subtypes, namely Gα_s_ and Gα_olf_, can be activated differentially with biased D1R agonists.^28^ In terms of CNS distribution, there is a considerable contrast in the localization of the two. Specifically, Gα_olf_ is highly segregated and enriched in the striatal regions while Gα_s_ is largely missing in the striatum and expressed in other areas such as the cortex.^29^ Selective activation of either Gα_s_ or Gα_olf_ *via* D1R has neuromodulatory effects in the cortex or striatum respectively.^28^ In fact, selectively activating D1R in the medial prefrontal cortex and the striatum has been suggested to be beneficial in treating the negative symptoms of schizophrenia and PD, respectively.^30^ In this regard, D1R compounds might have the advantage of achieving region-specific pharmacology. More evidence supports the idea that receptors can assume different active conformations to exert varying degrees of action at different transducers mediated by the same receptor.^31–33^ Furthermore, the G protein subtype selectivity profiles appeared to be translatable to the electrophysiological events and behavior of tested animals.^28, 34, 35^ Here, we characterize select D1R agonists for their transducer activation properties in cells, brain slices, and behavior. Our findings shed light on the conformational differences of D1R that transduce the activation of these two G proteins and the potential utility of separate targeting.

## RESULTS

### D1R agonists with differential efficacy at Gα_s_ or Gα_olf_ were identified in D1R-G protein engagement and activation assays

We used receptor-Gα engagement bioluminescence resonance energy transfer (BRET), Gα-γ activation BRET, and the fluorescence-based cAMP biosensor Pink Flamindo to study three main stages of activation: (1) Conformational change within the receptor to accommodate the G protein upon agonist binding, (2) subunit rearrangement/dissociation of heterotrimeric G protein following the GTP-GDP exchange, and (3) activation of immediate signaling effector.

Inference about the full or partial agonism of one ligand was made based on a reference compound, usually the endogenous ligand of the receptor studied. For all cell-based assays, DA was used as the reference and the efficacy of all ligands tested was normalized to that of DA in the same condition. In addition, SKF 81297 and SKF 38393, the established full and partial D1R agonists, respectively, were used to validate the assays’ sensitivity. As expected, SKF 81297 was as effective as DA in recruiting and activating both G proteins while SKF 38393 demonstrated significantly lower efficacy in both BRET configurations used **(Figure 1A, 1D, 2A, 2D).**

**Figure 1.**
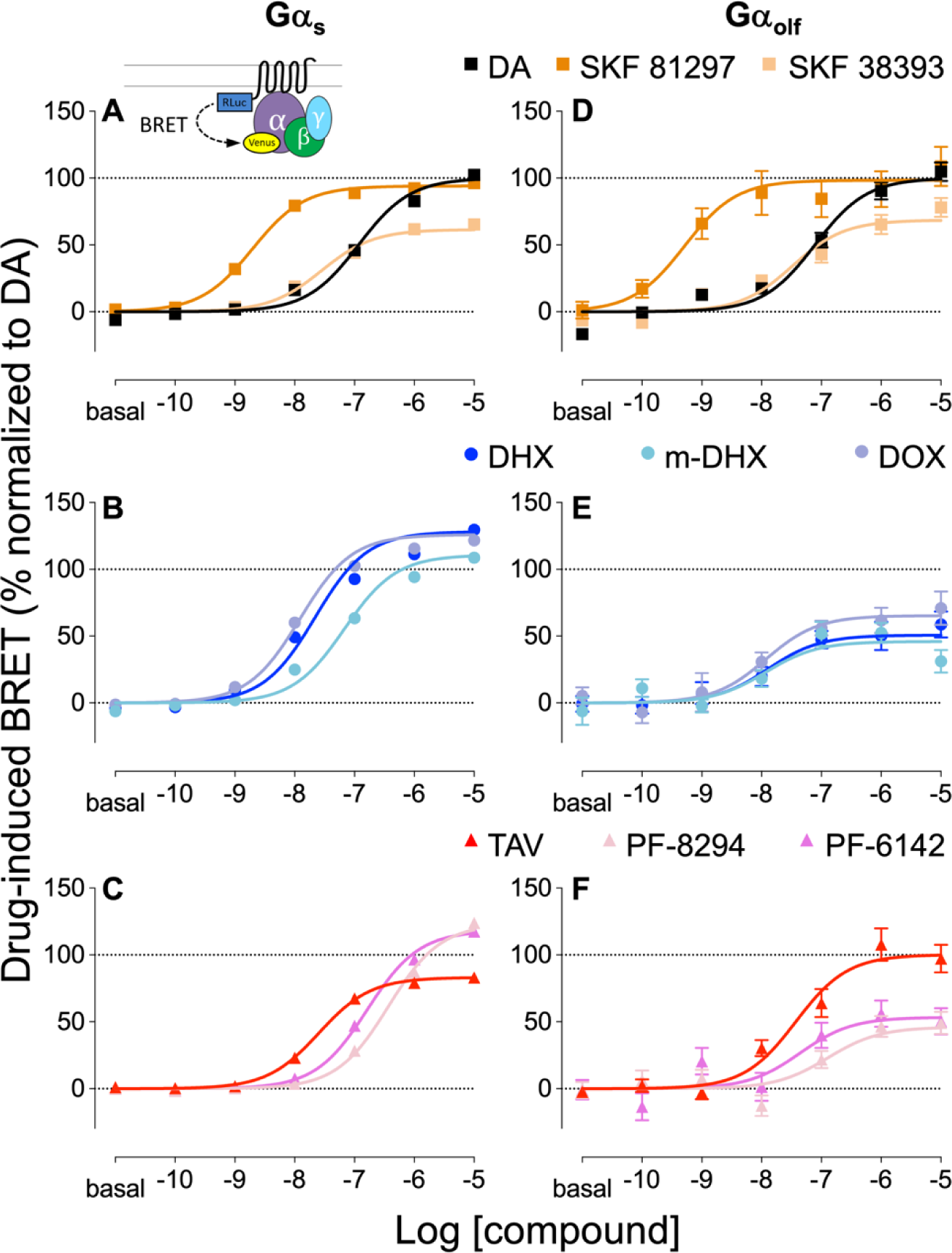
D1R agonist-induced D1R-Gα_s_ and -Gα_olf_ engagement. Drug-induced BRET ratio change between D1R-Rluc8 and Gα_s_-Venus **(A, B, C)** or Gα_olf_-Venus **(D, E, F)** in response to DA (black), SKF 81297 (dark orange), SKF 38393 (light orange) (top row), DHX (blue), m-DHX (light blue), DOX (violet) (middle row), TAV (red), PF-8294 (pink), and PF-6142 (magenta). Results were normalized to the E_max_ of DA response for D1R-Gα_s_ (**A, B, C**) and D1R-Gα_olf_ (**D, E, F**) and presented as mean ± SEM (n ≥ 5)

In this study, we investigated a series of D1R agonists with diverse structures from both catechol (dihydrexidine (DHX), methyl-dihydrexidine (m-DHX), doxanthrine (DOX))^36^ and non-catechol (tavapadon (TAV), PF-8294, PF-6142)^19^ classes. Unlike the control compounds (i.e., DA, SKF38393, SKF81297), there was a discernible difference in the efficacy of these agonists, relative to DA, at Gα_s_ and Gα_olf_. Of the compounds tested, DHX, m-DHX, DOX, PF-8294, and PF-6142 exerted full agonism at recruiting Gα_s_, evidenced by their comparable or significantly higher efficacy at Gαs, but acted as partial agonists in Gα_olf_ engagement. Interestingly, TAV induced Gα_olf_ coupling with comparable efficacy as DA but was significantly less efficacious at Gα_s_ recruitment **(Figure 1 B-F, Supplementary Table 1)**. Consistent with the results obtained from engagement BRET, the efficacy and potency of these agonists were retained when they were tested in activation BRET. TAV remained the only compound that behaved as a full agonist at Gα_olf_ activation while the rest of the compounds showed a clear preference for activating Gα_s_ over Gα_olf_.

### Gα_s_–Gα_olf_ subtype selectivity was consistently seen in G protein function at cAMP production level

Activation of adenylyl cyclase (AC) *via* Gα_s_ and Gα_olf_ to increase cAMP is the major signaling pathway mediated by D1R. Thus, drug-induced cAMP production was studied to evaluate Gα_s_ and Gα_olf_ activities at the functional level. To monitor the intracellular cAMP dynamic, we used the red-shifted fluorescence-based biosensor Pink Flamindo.^37^

Both full and partial benzazepine catechol agonists (SKF81297 and 38393) showed a consistent efficacy level in promoting cAMP production relative to DA as the G protein engagement or activation assays **(Figure 1A, 1D, 2A, 2D, 3A, 3D)**. The tetracyclic catechol compounds were equally efficacious as DA with Gα_s_ but were significantly less effective with Gα_olf_ **(Figure 3B, 3E)**. Similar to the G protein engagement and activation assays, PF-8294 and PF-6142 showed full and partial agonism in Gα_s_- and Gα_olf_-mediated cAMP production, respectively, while TAV behaved as a partial agonist at Gα_s_ and a full agonist at Gα_olf_ **(Figure 3C, 3F, Supplementary Table 3)**. Taken together, our assays have presented a distinct profile in G-protein activation of DHX, m-DHX, DOX, TAV, PF-8294 and PF-6142 at two highly homologous transducers, Gα_s_ and Gα_olf_, suggesting the possibility of developing compounds that selectively target one subtype.

**Figure 2.**
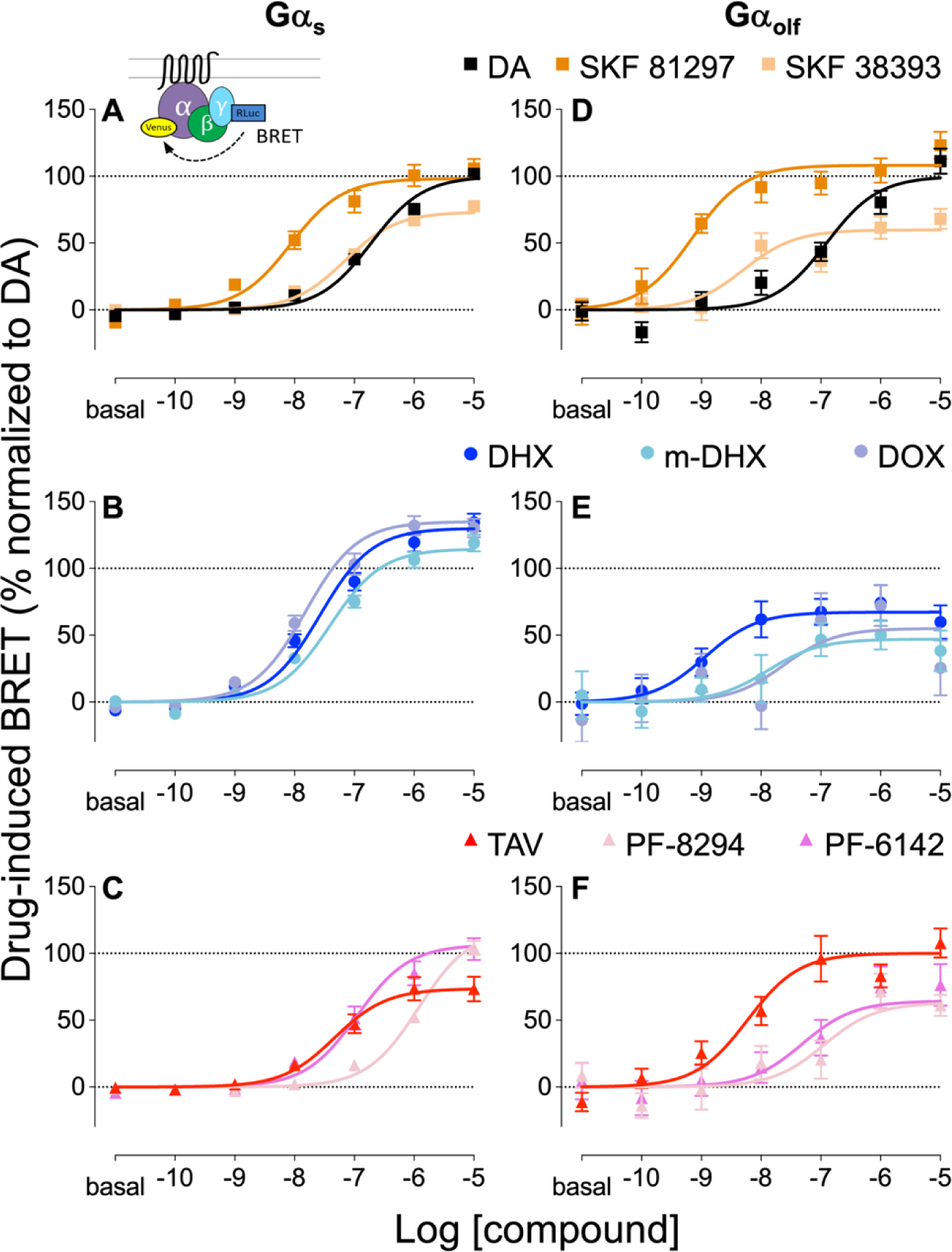
D1R agonist-induced D1R-Gα_s_ and -Gα_olf_ activation. Drug-induced BRET ratio change between γ7-Rluc8 and Gα_s_-Venus **(A, B, C)** or Gα_olf_-Venus **(D, E, F)** in response to DA (black), SKF 81297 (dark orange), SKF 38393 (light orange) (top row), DHX (blue), m-DHX (light blue), DOX (violet) (middle row), TAV (red), PF-8294 (pink), and PF-6142 (magenta). Results were normalized to the E_max_ of DA response for D1R-Gα_s_ (**A, B, C**) and D1R-Gα_olf_ (**D, E, F**) and presented as mean ± SEM (n ≥ 5)

**Figure 3.**
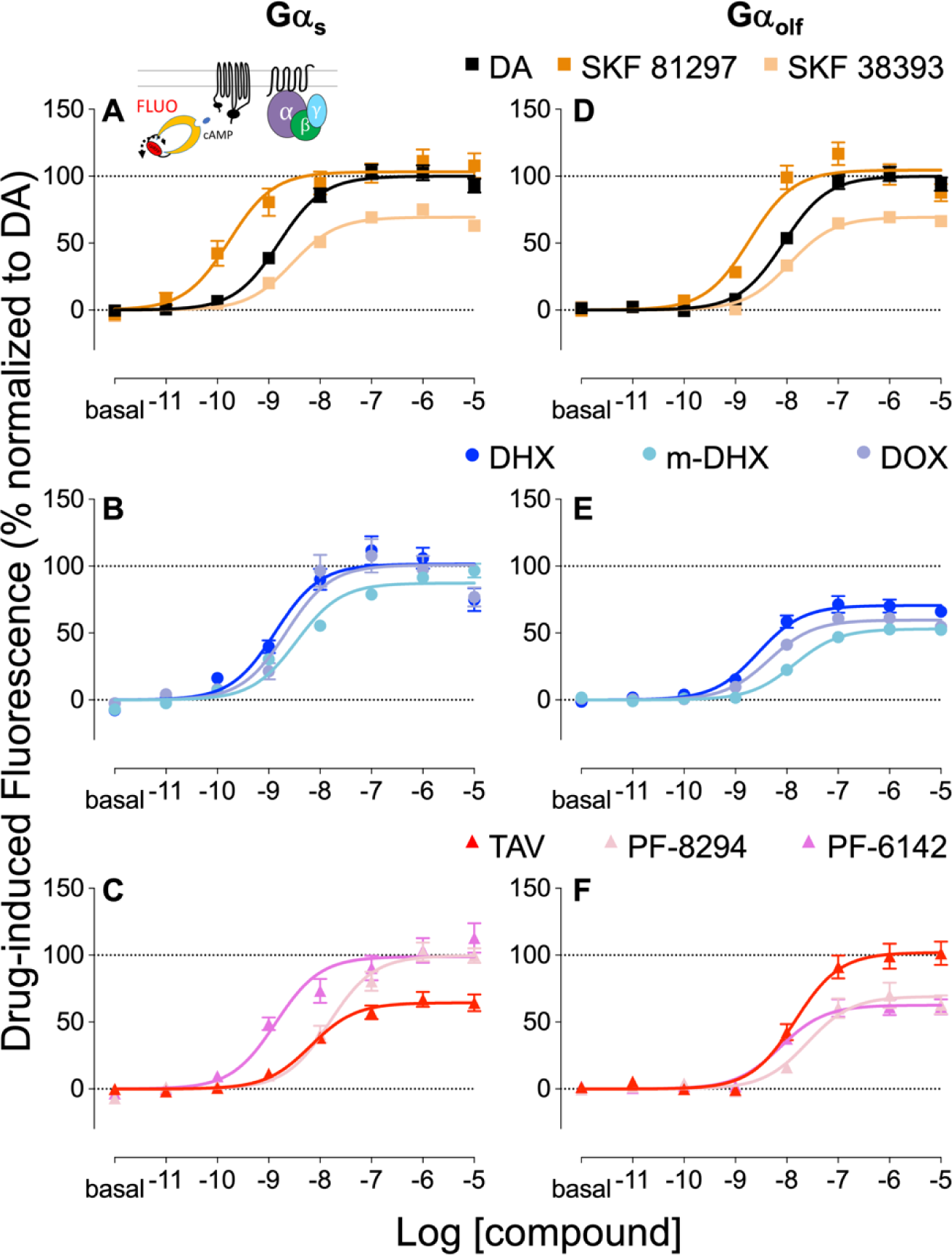
D1R agonist-induced cAMP production via Gα_s_ and Gα_olf_. Drug-induced fluorescence change of the cAMP biosensor Pink Flamindo in D1R-Gα_s_ **(A, B, C)** or D1R-Gα_olf_ **(D, E, F)** in response to DA (black), SKF 81297 (dark orange), SKF 38393 (light orange) (top row), DHX (blue), m-DHX (light blue), DOX (violet) (middle row), TAV (red), PF-8294 (pink), and PF-6142 (magenta). Results were normalized to the E_max_ of DA response for D1R-Gα_s_ (**A, B, C**) and D1R-Gα_olf_ (**D, E, F**) and presented as mean ± SEM (n ≥ 5)

### Pharmacological profiles for β-arrestin 1 and 2 recruitment were distinct from G protein engagement

β-arrestin-mediated GPCR internalization is a well-characterized major signaling event.^38^ In addition, β-arrestin also acts as a scaffold to recruit components of other signaling pathways independent of G-proteins.^39^ We used a BRET assay to study β-arrestin recruitment by these D1R agonists. As β-arrestin 1 and β-arrestin 2 are widely expressed in the CNS, we evaluated their activity at both subtypes relative to DA. SKF 38393 showed minimal recruitment of both β-arrestin 1 and 2 (22% and 28%, **Supplementary Table 4**). SKF 81297 was significantly less efficacious than DA in recruiting β-arrestin 1 and 2 (44% and 82%, **Supplementary Figure 1A, 1D, Supplementary Table 4**). Thus, notably, these benzazepine catechol compounds showed G-protein bias as reported previously.^40^

With the exception of m-DHX at β-arrestin 1 recruitment (70%), the tetracyclic catechol compounds demonstrated a high level of β-arrestin recruitment for both subtypes. Compared to their efficacy at Gα_s_ engagement BRET assay, DHX, m-DHX, and DOX appeared to be balanced agonists at these two transducers **(Supplementary Figure 1B, 1E)**. In contrast, the efficacy of the non-catechol compounds at β-arrestin 1 and 2 recruitment was significantly lower than that of DA. Cross-comparison with their E_max_ values at Gα_s_ coupling revealed a clear bias for G-protein engagement of TAV, PF-8294, and PF-6142.

### αN swap in Gα_olf_ chimera partially mimicked Gα_s_ activity

Recent cryo-EM structure studies suggested the receptor-G-protein interacting domains that contribute to the G protein subtype selectivity among different dopamine receptors.^41–43^ Of the domains involved at the interface, the intracellular loop 2 (ICL2) of D1R appeared to play a crucial role in this selectivity and was within interacting distance with the αN domain of the G protein.^20^ Previously, by substituting 6 amino acids in the αN helix of Gα_olf_ (ERLAYK) with the Gα_s_ equivalent residues (DKQVYR), referred to as Gα_olf_ αN, we were able to partially restore the Gα_s_ selectivity of DHX in engagement BRET.^28^ Here, we used the untagged Gα_olf/s_ αN and monitored the effect of this substitution at the functional level. Compared to their effects in wildtype (WT) Gα_s_ and Gα_olf_, SKF 81297 and SKF 38393 retained their relative efficacy and potency in cAMP production in Gα_olf_ αN **(Figure 4A)**. However, there is a clear difference in the efficacy of the other compounds when tested with the αN chimera. The tetracyclic catechol compounds (DHX, m-DHX, and DOX) significantly increased the cAMP production compared to their performance in WT Gα_olf_, approaching the level seen in Gα_s_ **(Figure 4B, Supplementary Figure 2B)**.

**Figure 4.**
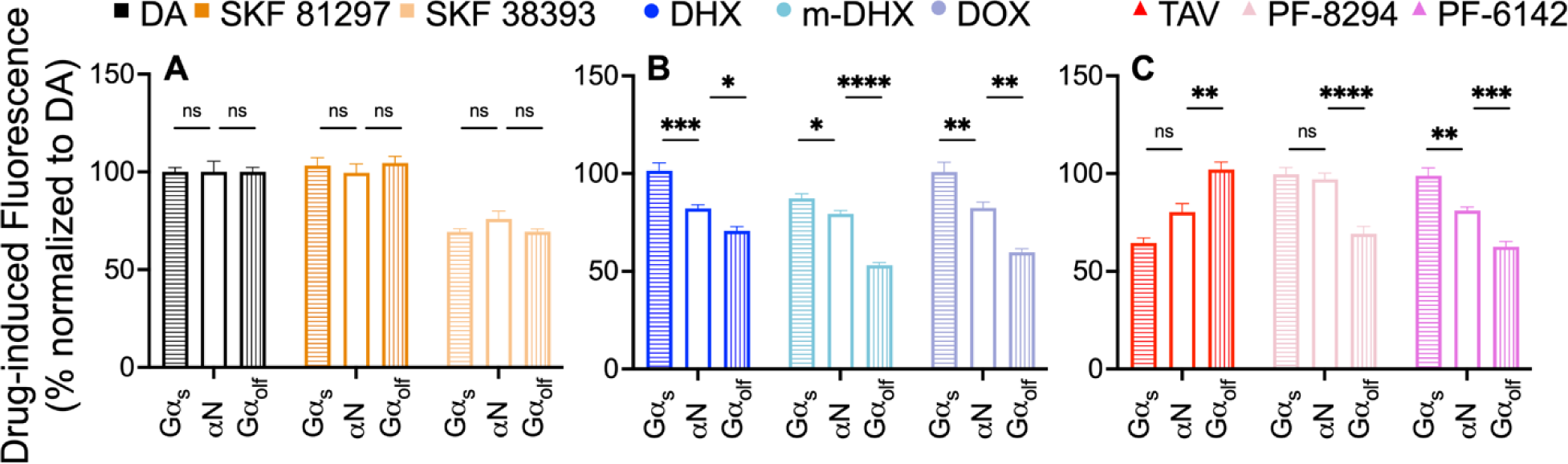
D1R agonist-induced cAMP production via Gα_olf/s_ chimera. Drug-induced fluorescence change of Pink Flamindo detecting cAMP level. **A**. DA (black), SKF81297 (orange), SKF38393 (light orange), **B**. DHX (blue), m-DHX (light blue), DOX (violet), **C**. TAV (red), PF-8294 (pink), and PF-6142 (magenta). Results were normalized to the E_max_ of DA response for Gα_s_, Gα_olf/s_ chimera (αN), Gα_olf_ and presented as mean ± SEM (n ≥ 5). Comparison of the E_max_ across Gα_s_, αN, and Gα_olf_ was done using one-way ANOVA followed by post-hoc Tukey test. *\*, **, ***, ****p<0.05, 0.01, 0.001, 0.0001,* respectively.

The effect of this substitution appeared to be more profound in the non-catechol compounds. We noticed the same trend in PF-8294 and PF-6142 where the efficacy in Gα_olf_ αN was significantly enhanced from their E_max_ in WT Gα_olf_. Strikingly, PF-8294 efficacy in Gα_olf_ αN was comparable to that in WT Gα_s_. TAV was the only Gα_olf_-selective compound in this set. When tested with Gα_olf_ αN, TAV was significantly less efficacious in producing cAMP than it was in WT Gα_olf_. In addition, TAV efficacy in Gα_olf_ αN was not different from that of Gα_s_ **(Figure 4C, Supplementary Figure 2C)**.

### D1R-mediated enhancement of pathway-specific synaptic events was seen with varied efficacy levels

Based on the distinct distribution of Gα_s_ and Gα_olf_ in the CNS, Gα_s_-selective D1R agonists are expected to exert higher efficacy in modulating cortex-related electrophysiological events whereas compounds that induce better Gα_olf_ coupling may preferentially modulate events that are associated with the striatum. The thalamus is one of the major sources of glutamate input to the cortex whereby it triggers cortical excitatory post-synaptic current (EPSC) via N-methyl-D-aspartate receptors (NMDAR). As D1R has been reported to potentiate the EPSC in the mPFC, recording EPSCs from the cortical pyramidal cells in the presence of D1R agonists may reveal NMDAR potentiation by D1R.^44–46^ On the other hand, activation of D1Rs at either axons or synaptic terminals of striatal medium spiny neurons (MSN) has been reported to presynaptically facilitate the SNr inhibitory post-synaptic current (IPSC).^47–49^ Thus, the IPSC recording of D1R-expressing MSN afferents in the SNr can evaluate D1R pharmacology with the degree of enhanced response.^50, 51^

Here, we combined optical stimulation of channelrhodopsin-2 (ChR2)^52^ and whole-cell voltage clamp on mouse brain slices to investigate D1R agonists’ effects on synaptic transmission in the thalamocortical and striatonigral projections specifically. AAV5-hSyn-hChR2(H134R)-EYFP was stereotaxically injected into the mediodorsal thalamus of *Drd1a-* tdTomato BAC transgenic mice for the expression in the thalamocortical pathway. In turn, the virus was injected into the dorsal lateral striatum of WT mice for the striatonigral pathway. Three weeks post-surgery, ChR2 was successfully expressed, evidenced by the strong expression of yellow fluorescence in the cortex **(Figure 5A)** or the SNr **(Figure 5D)**. D1R-expressing pyramidal cells in mPFC elicited an EPSC upon blue LED light stimulation. When the slices were perfused with 1 μM SKF 81297, the EPSC peak amplitude significantly increased (113.4% of baseline, p<0.01). The same results were observed when the slices were perfused with 1 μM DHX (113.1% of baseline, p<0.05). However, perfusing 1 μM tavapadon did not enhance the cortical EPSC amplitude **(Figure 5B, 5C)**.

**Figure 5.**
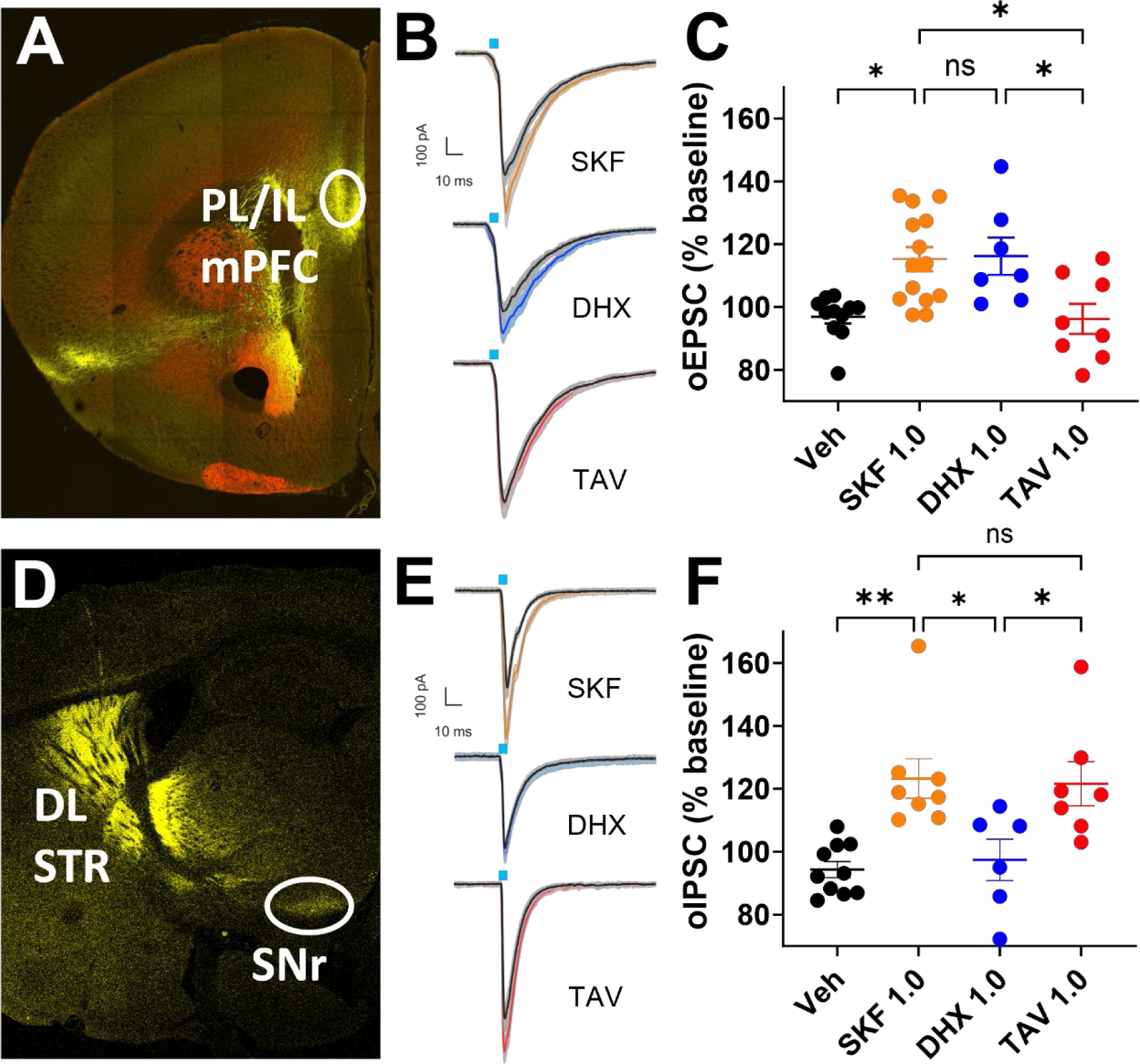
Gα_s_ and Gα_olf_ activities in brain slice electrophysiology. ChR2-YFP injection in the mediodorsal thalamus projecting to the prelimbic/infralimbic mPFC of D1R-tdTomato mouse **(A)** and in the dorsal striatum projecting to the SNr of WT mouse **(D)**. Average traces of optically-elicited ChR2-driven EPSCs in the mPFC **(B)** and IPSCs in the SNr **(E)** following 2 ms of photostimulation by blue light. Analysis of the D1R-mediated potentiation of EPSCs in the mPFC **(C)** IPSCs in the SNr **(F)** was done using one-way ANOVA followed by post-hoc Tukey test. *, **p<0.05, 0.01, respectively

Photostimulation of the MSN GABAergic afferent in the SNr elicited an IPSC that was markedly enhanced by 1 μM SKF 81297 (123.3% of baseline, p<0.01). In contrast to its effect in the mPFC, 1 μM tavapadon significantly potentiated the IPSC in the SNr (123.1% of baseline, p<0.05). This effect was comparable to that seen with SKF 81297. DHX, on the other hand, failed to modulate the IPSC in the SNr **(Figure 5E, 5F)**. The electrophysiological effects seen from the recordings with DHX and TAV were consistent with their selectivity profiles *in* vitro **(Table 1, 2, 3)**. This suggested that D1R agonists that preferentially activate one G protein subtype might be able to selectively modulate the electrophysiological events in the area where that G protein is enriched, indicating that D1R region-specific pharmacology can be achieved by targeting the G-protein subtypes.

### Locomotor rescue in cataleptic mice showed a significant difference in efficacy between tavapadon and dihydrexidine

D1R in the striatum regulates locomotor activity, likely via Gα_olf,_ due to its high level of expression in this brain region.^53^ The psychostimulatory effect of D1R agonists reflects this Gα_olf_-mediated striatal function. However, endogenous DA can be a potential confounding factor in studying this effect *in vivo* as it can mask the effects of D1R agonists via D1R in both the striatum and elsewhere. Therefore, it is necessary to deplete the endogenous DA levels to elicit akinetic state so that D1R agonist effects in the striatal neurons can be studied effectively as widely reported for DA drugs. ^54–56^ To that end, reserpine was used to irreversibly block the vesicular monoamine transporters 1 and 2 (VMAT-1 and VMAT-2), rendering the depletion of DA in the synaptic terminals.^57^

SKF 81297, DHX, and TAV induced significant locomotor stimulation in a dose-dependent manner **(Figure 6A, 6B, 6C)**. At the highest tested dose (6 mg/kg), the effects of TAV and SKF 81297 were comparable. Interestingly, DHX was significantly less efficacious than either of the aforementioned drugs in stimulating locomotor activity (44.79% of SKF 81297 and 45.09% of TAV) **(Figure 6D)**. This is in line with our *in vitro* and *ex vivo* data and suggests that Gα_olf_-selective D1R agonists might be better at rescuing locomotor activity, probably due to their preferential stimulation of D1Rs in the striatum.

**Figure 6.**
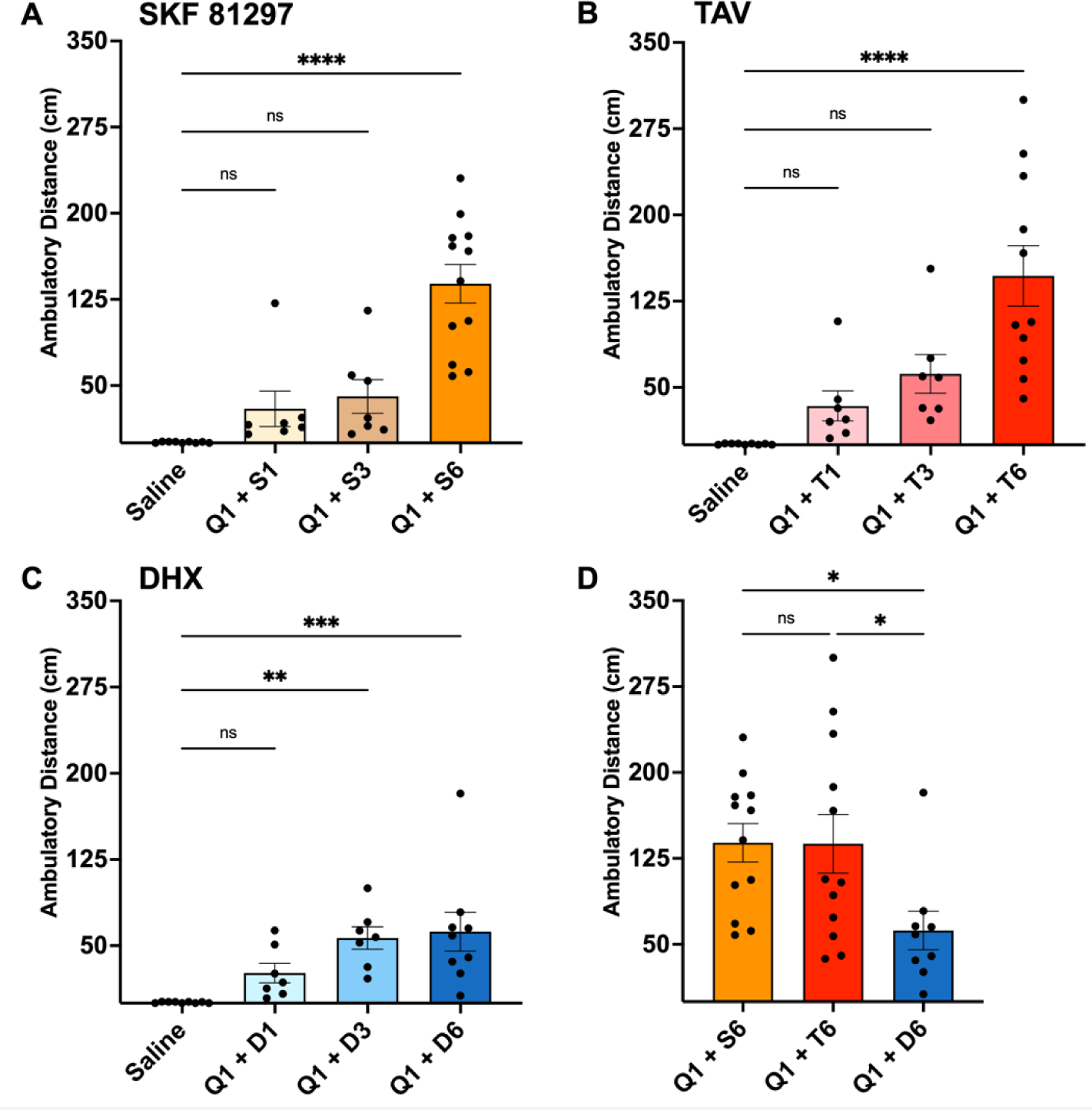
D1R agonists effect on locomotor activity. Psychomotor stimulatory effect of SKF-81297 **(A)**, TAV **(B),** and DHX **(C)** on DA-depleted mice. *, **, ***, **** p<0.05, 0.01, 0.001, 0.0001, respectively, compared to the effect of saline using one-way ANOVA followed by post-hoc Dunnett test. **D.** Comparison of the ambulatory distance induced by these drugs at 6 mg/kg was done using one-way ANOVA followed by post-hoc Tukey test. *, **, ***, **** p<0.05, 0.01, 0.001, 0.0001, respectively

## DISCUSSION

It has been reported that the restricted conformation of the β-phenyl-dopamine pharmacophore renders the selectivity and full agonism at D1R of catechol D1R agonists.^36^ Here, we expanded on this finding and revealed a differential activation profile of DHX, m-DHX, and DOX at two homologous G protein subtypes, Gα_s_ and Gα_olf_. While these compounds exerted full agonism at Gα_s_, they behaved as partial agonists at Gα_olf_. This characteristic was discernibly different from that of the benzazepine catechol D1R agonists, where the β-phenyl ring is not restricted. We surmised the rigid ring structure of these compounds might contribute to the activity bias towards Gα_s_.

A series of arylphenoxyaryl D1R agonists was developed in recent years and their structure-activity relationship has been studied by several groups. Modifications of the two heterocyclic groups at the ends or the core linking structure could drastically alter the selectivity of these compounds between G protein and β-arrestin signaling.^18, 19, 26, 27^ Replacing the trifluoromethyl pyridine ring of TAV with the pyridofuran ring of PF-8294 and PF-6142 increased the potency in the cAMP production assay and also enhanced both efficacy and potency in the β-arrestin recruitment assay.^26^ On the other hand, substituting the imidazopyridine ring of PF-8294 with the imidazopyrazine ring of PF-6142 at a different attachment position increased the efficacy and potency in the cAMP production assay, but did not cause a significant change in the β-arrestin recruitment assay.^19, 27^ These findings align with our current data on Gα_s_-mediated cAMP production and β-arrestin recruitment (Figure 3C, Supplementary Figure 1). To our knowledge, we are the first to report the profiles of these compounds at Gα_s_ and Gα_olf_. Interestingly, TAV, PF-8294, and PF-6142 consistently showed differential activities at these two G proteins. While PF-8294 and PF-6142 demonstrated selectivity toward Gα_s_, TAV was selective toward Gα_olf_. This suggested that by substituting the heterocyclic groups of the arylphenoxyaryl pharmacophore, in addition to altering the G protein - β-arrestin relationship, subtype selectivity can also be modified. This is significant given the unique expression pattern of Gα_s_ and Gα_olf_ in the CNS and the behaviors associated with these regions.

The β-arrestin-mediated GPCR internalization accounts for developing drug tolerance.^58^ The high efficacy at β-arrestin recruitment of DHX, m-DHX, and DOX might explain the rapid tolerance usually seen with tetracyclic catechol D1R agonists.^16^ Our results on β-arrestin recruitment of PF-8294 and PF-6142 agreed with the previously published data regarding the reduced desensitization of D1R upon drug exposure.^27^ Furthermore, the low β-arrestin recruitment of TAV could be one reason for this compound’s improved tolerability in clinical trials.^23^

Our previous publication showed that the αN/IL2 (Gα/D1R) interface might contribute to the G protein subtype selectivity of these compounds.^28^ While the chimeric construct significantly altered the efficacy of these compounds trending toward Gα_s_, their potency remained unchanged from WT Gα_olf_ (Supplementary Table 3), suggesting that other domains might also contribute to the G protein subtype selectivity. Despite the high degree of homology, there are domains with divergent amino acid sequences between Gα_s_ and Gα_olf_ especially at the regions interfacing D1R. More thorough analysis of subdomains is warranted and currently underway.

Our electrophysiology data indicated a pathway-specific effect of DHX and TAV. We expected that selective targeting of G protein subtype via D1R would allow modulation of the neurological and neuropsychiatric outcomes associated with that region. Specifically, cortical D1R has a significant role in regulating executive functions and working memory, ^8, 10, 59^ while striatal D1R is one of the main players in voluntary movements, reward, and motivation.^4, 5, 7, 60–62^ Thus, region-specific D1R activation will have great implications for developing treatments for several disorders, such as cognitive deficiency and movement disorder, while potentially limiting the adverse off-target effects from globally activating all the D1R in the brain.

Developing D1R agonists, especially ones with the catechol moiety, has been proven to be challenging due to appreciable adverse effects. It has been reported that some compounds are epileptogenic or can decrease seizure thresholds.^63, 64^ In addition, cardiovascular effects are the other concern regarding the safety of these agonists. In fact, hypotension prevented several compounds from advancing to later phases of clinical trials,^14, 16^ which might be due to the off-target effects of these drugs on peripheral D1Rs.^65^ In fact, some degrees of seizures were observed in a few mice when tested at 6 mg/kg of SKF 81297 although this phenomenon was not observed with DHX and especially Gα_olf_-biased TAV, even at 10 mg/kg (data not shown).

Based on our *in vitro* and electrophysiology results, we expected that TAV might impact striatum-mediated locomotor activity to a higher degree than DHX. Indeed, our results on locomotor rescue of these D1R agonists showed a significantly higher efficacy of TAV compared to DHX **(Figure 6D)**. This finding was particularly interesting because it corroborated our hypothesis that it would be possible to selectively activate D1R in different brain regions by exploiting the unique expression pattern of Gα_s_ and Gα_olf_ in the CNS.

Overall, our study suggested that the *in vitro* G protein subtype selectivity profile of D1R agonists is translatable to their physiological effects and that the current design is a reliable model to screen for compounds with different clinical significance. The results reported at various physiological levels herein consistently showed that TAV had a unique efficacy bias towards Gα_olf_ while PF-8294, PF-6142 and the DHX analogs series showed a Gα_s_ efficacy bias.

## SIGNIFICANCE

Functional selectivity of transducer-coupling in G protein-coupled receptors is a promising concept in drug development that has led to targeted signaling approaches. While functional selectivity between G proteins and β-arrestins has opened up various avenues for therapeutics, mechanistic understanding of G protein subtype selectivity, in particular, within the same G protein subfamilies lags far behind. Here, we report a first case of Gα_olf_-biased agonist (tavapadon) in the dopamine D1 receptor. Together with Gα_s_-biased agonist (dihydrexidine), Gα_s_ vs. Gα_olf_ subtype selectivity within the same Gα_s_ subfamily was first evaluated at discrete configurations via bioluminescence resonance energy transfer assays. Collectively, they detected D1 receptor engagement, G protein activation, and adenylyl cyclase activation of specific Gα subunit. One of the interacting domains that distinguish the two subtypes was validated by a chimeric mutant containing a swapped domain. Gα_s_- and Gα_olf_-mediated D1 receptor effects via pathway-specific electrophysiology verified the biased nature of these two ligands in transgenic mice. Lastly, Gα_olf_-mediated D1 receptor activation in the striatum in the reserpinized mouse model allowed for functional validation of G_olf_-biased agonist effect at the behavioral level. Taken together, the current study demonstrates the first case of biased pharmacology for two subtypes of Gαs subfamily through various physiological levels. This proof-of-concept work provides possibilities for cell-type-specific pharmacology as Gα expression pattern can be exploited, in this case, in the brain. In the treatment of movement disorders, Gα_olf_-biased D1 receptor agonists may have more desired outcomes by circumventing adverse effects from Gα_s_ activation.

## MATERIALS AND METHODS

### Cell-based Assays

#### Transfection

WT or Gα_s_- and Gα_olf_-KO HEK 293T cells (CL4) were seeded at 4 million cells per 10-cm plate and cultured in Dulbecco’s modified Eagle’s medium (DMEM) containing 10 % fetal bovine serum (FBS), 2 mM L-glutamine and 1 % penicillin-streptomycin at 37 °C and incubated in 5 % CO_2_ – 95 % moisturized air. A constant amount of cDNA was transfected using polyethyleneimine (PEI) (1:2 w/w). For engagement BRET assays, CL4 cells were transfected with D1R-RLuc (Renilla Luciferase), Gα_ss_154-Venus or Gα_olf_155-Venus, β2, γ7, and RIC8B. For activation BRET assays, CL4 cells were transfected with untagged D1R, Gα_ss_67-Venus or Gα_olf_69-Venus, β2, γ7-L8, and RIC8B. For cAMP production assays, CL4 cells were transfected with D1R, Gα_ss_ or Gα_olf_ or Gα_olf_ αN, β2, γ7, RIC8B, and Pink Flamindo (a fluorescence-based cAMP biosensor). For β-Arrestin recruitment assays, WT cells were transfected with D1R-RLuc, β-Arrestin 1-Venus or β-Arrestin 2-Venus, and GRK2. The ratio between the transfected constructs had been previously optimized for the dynamic range of the drug-induced BRET change as described previously.^28^

#### Bioluminescence resonance energy transfer (BRET)

Forty-eight hours post-transfection, cells were harvested and resuspended in potassium-based buffer (140 mM KCl, 10 mM NaCl, 1 mM MgCl_2_, 0.1 mM KEGTA, 20 mM NaHEPES, pH 7.2) for engagement and activation BRET or phosphate-buffered saline (PBS) containing 200 μM sodium bisulfite (NaBi) for β-Arrestin recruitment assays. Cells were distributed at 100,000 cells per well in a white 96-well plate and 5 μM coelenterazine H was added. Ligands at varying concentrations were added 1 minute later. For experiments with Gα_olf_155-Venus and Gα_olf_69-Venus, cells were pre-incubated with 10 μg/ml digitonin for 60 minutes prior to ligand addition. Bioluminescence and fluorescence emissions were recorded at 485 and 530 nm, respectively.

#### cAMP production assay

Cells were prepared as described above for the BRET assay, except that they were resuspended in PBS containing 200 μM NaBi and distributed in a black 96-well plate. Plate reader is set at 540 nm excitation and 590 nm emission. For all *in vitro* experiments, dose-response curves and data analysis were done using GraphPad Prism10.

### Mouse-based Assays

#### Stereotaxic surgery

*Drd1a-*tdTomato BAC transgenic mice were anesthetized and maintained with isoflurane/oxygen through a nose cone mounted on a stereotaxic apparatus. The scalp was opened, and two holes were drilled in the skull (-0.4 mm AP, ±0.4 mm ML, -3.5 mm DV from bregma for the thalamocortical pathway; 0 mm AP, ±2 mm ML, -3 mm DV from bregma for the nigrostriatal pathway). AP, ML, DV, which stands for anterior-posterior, medial-lateral, and dorsal-ventral, respectively, were the spatial coordinates relative to the bregma. AAV5-hSynapsin-hChannelrhodopsin 2(H134R)-EYFP virus at titer of 2 x 10^13^ vg/ml (0.5 μl/side) was injected at 0.1 μl/min. The needle was left in place for 1 minute before being slowly removed.

#### Slice electrophysiology

Horizontal (for the nigrostriatal pathway) and coronal slices (for the thalamocortical pathway) at 220 µm thickness were prepared using a vibratome (VT-1000S, Leica). Mice were anesthetized and perfused with modified artificial cerebral spinal fluid (m-aCSF) containing (in mM): 92 NMDG, 20 HEPES, 25 glucose, 30 NaHCO_3_, 1.2 NaH_2_PO_4_, 2.5 KCl, 5 sodium ascorbate, 3 sodium pyruvate, 2 thiourea, 10 MgSO_4_, 0.5 CaCl_2_, 300–310 mOsm, and pH 7.3–7.4. Slices were sectioned in cold m-aCSF, recovered at 32 °C in the same buffer saturated with carbogen for 10 min, and transferred to a holding chamber filled with carbogen-saturated aCSF (holding aCSF) containing (in mM): 92 NaCl, 20 HEPES, 25 glucose, 30 NaHCO_3_, 1.2 NaH_2_PO_4_, 2.5 KCl, 5 sodium ascorbate, 3 sodium pyruvate, 2 thiourea, 1 MgSO_4_, 2 CaCl_2_, 300–310 mOsm, and pH 7.3–7.4. During recordings, slices were continuously perfused at 2 ml/min with carbogen-saturated aCSF containing (in mM): 124 NaCl, 2.5 KCl, 1.25 NaH_2_PO_4_, 1 MgCl_2_, 26 NaHCO_3_, 11 glucose, 2.4 CaCl_2_, 300–310 mOsm, and pH 7.3–7.4. The temperature of the recording chamber was maintained at 31–32 °C. Electrodes (3–5 MΩ) iwere backfilled with an internal solution containing (in mM): 120 mM K gluconate, 20 KCl, 0.05 EGTA, 10 HEPES, 1.5 MgCl_2_, 2.18 Na_2_ ATP, 0.38 Na GTP, 10.19 Na phosphocreatine, 280–285 mOsm, and pH 7.3–7.4. Pyramidal neurons in layer V of the mPFC and GABAergic neurons in the SNr were visually identified using IR-DIC optics with an upright microscope (Olympus). Whole-cell voltage-clamp recordings (held at resting membrane potential) were made using a MultiClamp 700B amplifier (2 kHz low-pass Bessel filter and 10 kHz digitization) with pClamp 11 software (Molecular Devices).

For ChR2 evoked responses, the thalamocortical and striatonigral terminals were activated using a 473 nm DPSS blue laser. The laser was controlled by an Optogenetics TTL Pulse Generator (Doric Lenses, Quebec, Canada). The stimulation protocol consisted of 4 pulsed lights at 5 s intervals with a 2 ms pulse duration at 1 mW, followed by a 40 s pause. After voltage-clamp recording was established, cell response to photostimulation in the absence of the drugs (baseline) was recorded for 20 minutes followed by 6 – 10 minutes of drug perfusion. Peak amplitudes in the last 3 minutes of each condition were used for statistical analysis.

For the thalamocortical pathway, the following numbers of slices were recorded for each drug condition: SKF 81297 (15 recordings from 7 animals), DHX (8 recordings from 3 animals), TAV (8 recordings from 6 animals). For the nigrostriatal pathway, the following numbers of slices were recorded for each drug condition: SKF 81297 (8 recordings from 4 animals), DHX (6 recordings from 5 animals), TAV (7 recordings from 5 animals). All data were reported as mean ± SEM. Data were analyzed in Clampex and statistical analysis was done using GraphPad Prism 10.

#### Locomotor assay

Reserpine 3 mg/kg (injection volume 10 μl/g bodyweight) was administered subcutaneously 18 – 20 h before undertaking the behavioral assays. Following the 18-20 h reserpine effect, mice were injected intra-peritoneally with a cocktail of quinpirole 1 mg/kg and either SKF 81297, DHX, or TAV at 1, 3, 6 mg/kg (injection volume 10 μl/g bodyweight). Mice were introduced to a 28-cm x 28-cm open-field arena (ENV-520, Med Associate Inc.) for a 60-minute recording session. Data was reported as the mean ambulatory distance (cm) in 15 minutes.

## AUTHOR CONTRIBUTIONS

HY designed the experiments. AMN, AS, and VQ carried out the experiments, and performed the data analysis with HY. AI provided Gα_s_- and Gα_olf_-KO HEK 293T cells. DEN provided tetracyclic catechol D1R agonists. AMN and HY prepared the initial manuscript, with contributions from all the authors in finalizing the manuscript.

## ACKNOWLEDGEMENTS

PF-8294 and PF-6142 compounds were provided under a Material Transfer Agreement with F. Hoffmann-La Roche LTD (Basel, Switzerland). The work was supported by Northeastern University Startup Funds and Tier1 Intramural Grant (HY).

