## Supporting information for "Characterization of Gα_s_ and Gα_olf_ activation by catechol and non-catechol dopamine D1 receptor agonists"

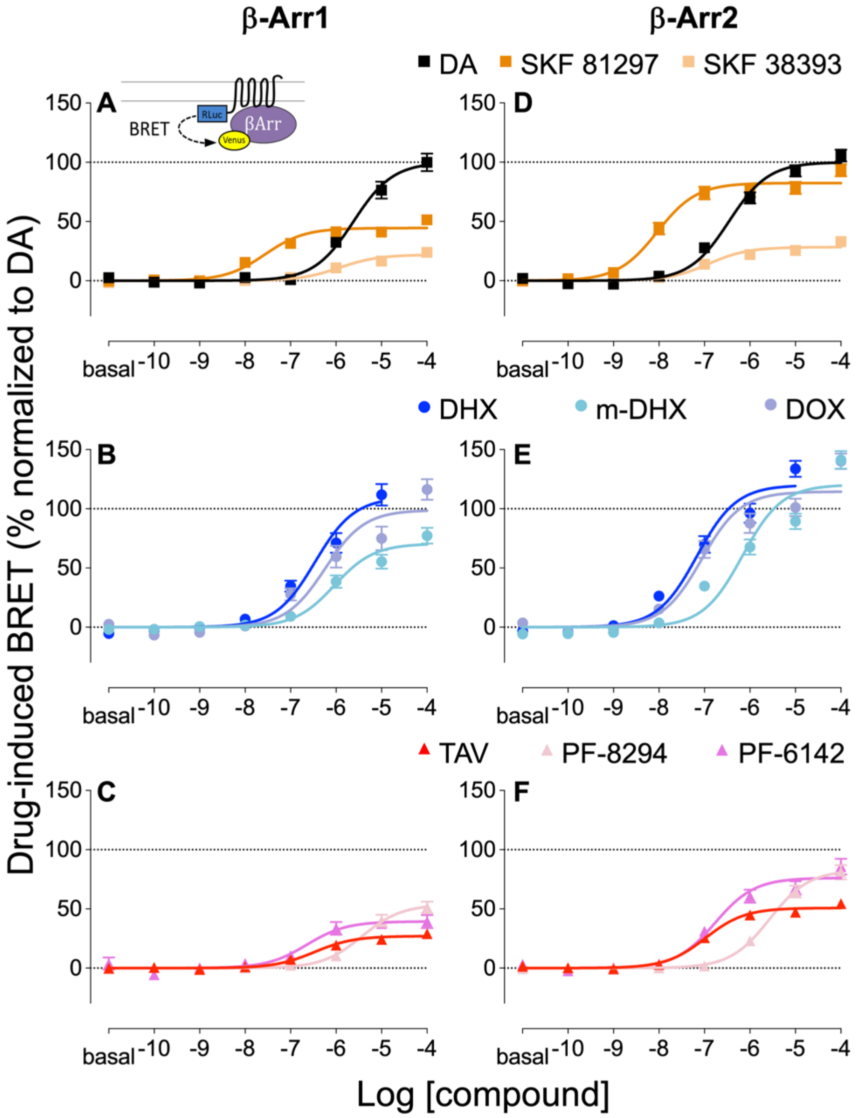


**Supplementary Figure 1. D1R agonist-induced D1R-β-arrestin 1 and β-arrestin 2 recruitment.** Drug-induced BRET ratio change between D1R-Rluc8 and β-arrestin 1 **(A, B, C)** or β-arrestin 2 **(D, E, F)** in response to DA (black), SKF 81297 (dark orange), SKF 38393 (light orange) (top row), DHX (blue), m-DHX (light blue), DOX (violet) (middle row), TAV (red), PF-8294 (pink), and PF-6142 (magenta). Results were normalized to the $E_{max}$ of DA response for D1R- β-arrestin 1 (**A, B, C**) and D1R- β-arrestin 2 (**D, E, F**) and presented as mean ± SEM (n ≥ 5)

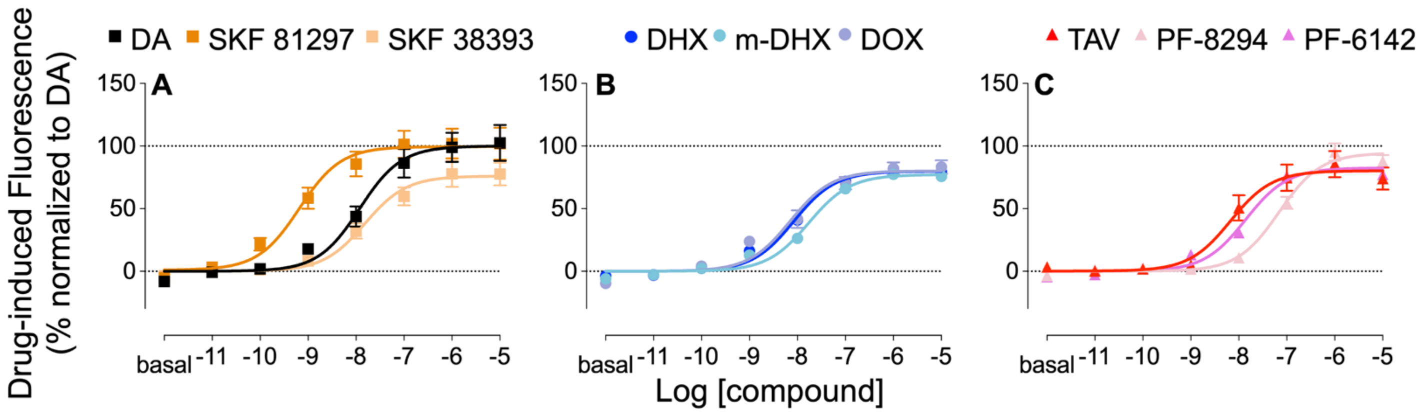


**Supplementary Figure 2. D1R agonist-induced cAMP production via G$\alpha_{olf/s}$ chimera.** Drug-induced fluorescence change of the cAMP biosensor Pink Flamindo in D1R-G$\alpha_{olf/s}$ in response to **A.** DA (black), SKF 81297 (dark orange), SKF 38393 (light orange), **B.** DHX (blue), m-DHX (light blue), DOX (violet), **C.** TAV (red), PF-8294 (pink), and PF-6142 (magenta). Results were normalized to the $E_{max}$ of DA response and presented as mean ± SEM ($n \geq 5$)

**Supplementary Table 1. Pharmacological comparison of $G\alpha_s$ and $G\alpha_{olf}$ engagement**

| | $G\alpha_s$ | | $G\alpha_{olf}$ | |
|---|---|---|---|---|
| | $E_{max}$ (%DA) | $EC_{50}$ (nM) | $E_{max}$ (%DA) | $EC_{50}$ (nM) |
| **DA** | 100.00 ± 1.02 | 130.92 ± 7.28 | 100.00 ± 3.16 | 78.16 ± 19.61 |
| **SKF 81297** | 96.46 ± 0.81 | 1.94 ± 0.14 | 98.40 ± 6.19 | 0.52 ± 0.54 |
| **SKF 38393** | 62.98 ± 0.57**** | 28.18 ± 2.14 | 68.51 ± 3.24**** | 32.89 ± 14.86 |
| **DHX** | 131.30 ± 1.70**** | 21.98 ± 2.37 | 50.70 ± 5.00**** | 12.79 ± 14.75 |
| **m-DHX** | 106.9 ± 1.415** | 67.76 ± 6.89 | 45.93 ± 4.97**** | 11.72 ± 13.39 |
| **DOX** | 129.00 ± 1.96**** | 12.27 ± 1.51 | 65.32 ± 5.69*** | 11.80 ± 9.7 |
| **TAV** | 83.29 ± 0.72**** | 25.35 ± 1.44 | 101.70 ± 5.39 | 43.55 ± 20.08 |
| **PF-8294** | 126.20 ± 1.58**** | 369.83 ± 31.11 | 46.03 ± 5.74**** | 127.64 ± 136.59 |
| **PF-6142** | 121.10 ± 1.50**** | 163.68 ± 14.22 | 53.38 ± 7.03**** | 44.16 ± 60.85 |

Data were presented as mean ± SEM. *, **, ***, *****p<0.05, 0.01, 0.001, 0.0001,* respectively, compared to DA using one-way ANOVA followed by post-hoc Dunnett test.

**Supplementary Table 2. Pharmacological comparison of $G\alpha_s$ and $G\alpha_{olf}$ activation**

| | $G\alpha_s$ | | $G\alpha_{olf}$ | |
|---|---|---|---|---|
| | $E_{max}$ (%DA) | $EC_{50}$ (nM) | $E_{max}$ (%DA) | $EC_{50}$ (nM) |
| **DA** | 100.00 ± 1.74 | 190.99 ± 23.10 | 100.00 ± 6.86 | 148.25 ± 67.32 |
| **SKF 81297** | 98.29 ± 3.99 | 7.64 ± 2.49 | 107.50 ± 4.96 | 0.89 ± 0.61 |
| **SKF 38393** | 73.28 ± 1.68**** | 69.20 ± 12.26 | 57.46 ± 4.47**** | 5.06 ± 4.99 |
| **DHX** | 130.10 ± 3.26**** | 26.67 ± 5.96 | 62.29 ± 5.52** | 1.01 ± 1.14 |
| **m-DHX** | 114.70 ± 2.93** | 39.26 ± 8.62 | 42.74 ± 8.12**** | 10.94 ± 30.81 |
| **DOX** | 135.00 ± 3.23**** | 15.17 ± 3.23 | 45.78 ± 12.16*** | 20.18 ± 98.89 |
| **TAV** | 73.60 ± 3.75**** | 46.67 ± 21.46 | 99.53 ± 5.51 | 6.98 ± 3.84 |
| **PF-8294** | 118.3 ± 3.29*** | 1188.50 ± 193.25**** | 52.22 ± 8.08*** | 72.11 ± 134.14 |
| **PF-6142** | 106.50 ± 4.14 | 110.92 ± 33.53 | 53.04 ± 7.92** | 27.35 ± 59.44 |

Data were presented as mean ± SEM. *, **, ***, ****$p<0.05$, *0.01, 0.001, 0.0001,* respectively, compared to DA using one-way ANOVA followed by post-hoc Dunnett test.

**Supplementary Table 3. Pharmacological comparison of cAMP production via $G\alpha_s$, $G\alpha_{olf}$, $G\alpha_{olf/s}\ \alpha N$**

| | $G\alpha_s$ | | $G\alpha_{olf}$ | | $G\alpha_{olf/s}\ \alpha N$ | |
|---|---|---|---|---|---|---|
| | $E_{max}$ (%DA) | $EC_{50}$ (nM) | $E_{max}$ (%DA) | $EC_{50}$ (nM) | $E_{max}$ (%DA) | $EC_{50}$ (nM) |
| **DA** | 100.00 ± 2.26 | 1.60 ± 0.34 | 100.00 ± 2.32 | 8.55 ± 1.55 | 100.00 ± 5.43 | 11.89 ± 5.41 |
| **SKF 81297** | 103.30 ± 3.93 | 0.19 ± 0.09 | 104.60 ± 3.37 | 1.89 ± 0.56 | 99.53 ± 4.59 | 0.66 ± 0.31 |
| **SKF 38393** | 69.36 ± 1.63$^{****}$ | 2.83 ± 0.58 | 69.40 ± 1.48$^{****}$ | 11.27 ± 1.71 | 76.01 ± 3.96$^{***}$ | 17.66 ± 4.25 |
| **DHX** | 101.50 ± 3.94 | 0.98 ± 0.23 | 70.68 ± 2.22$^{****}$ | 3.44 ± 0.96 | 82.17 ± 1.94$^{**}$ | 8.41 ± 1.50 |
| **m-DHX** | 87.32 ± 2.40 | 1.99 ± 0.35 | 53.12 ± 1.38$^{****}$ | 15.56 ± 3.16 | 79.43 ± 1.71$^{***}$ | 17.18 ± 2.98 |
| **DOX** | 100.80 ± 5.01 | 2.26 ± 1.06 | 59.72 ± 1.86$^{****}$ | 3.52 ± 0.97 | 82.47 ± 2.90$^{**}$ | 7.07 ± 2.00 |
| **TAV** | 64.45 ± 2.58$^{****}$ | 6.56 ± 2.20 | 102.00 ± 3.95 | 14.29 ± 4.24 | 80.25 ± 4.43$^{**}$ | 6.71 ± 3.26 |
| **PF-8294** | 99.67 ± 3.39 | 13.65 ± 3.74 | 69.21 ± 3.72$^{****}$ | 28.58 ± 10.08 | 97.10 ± 3.15 | 69.02 ± 14.13$^{****}$ |
| **PF-6142** | 98.89 ± 4.00 | 3.63 ± 1.64 | 62.64 ± 2.58$^{****}$ | 15.70 ± 9.79 | 80.99 ± 1.91$^{**}$ | 13.87 ± 2.33 |

Data were presented as mean ± SEM. **, **, ***, ****$p<0.05, 0.01, 0.001, 0.0001$,* respectively, compared to DA using one-way ANOVA followed by post-hoc Dunnett test.

**Supplementary Table 4. Pharmacological comparison of β-arrestin 1 and 2 recruitment**

| | β-arrestin1 | | β-arrestin2 | |
|---|---|---|---|---|
| | $E_{max}$ (%DA) | $EC_{50}$ (nM) | $E_{max}$ (%DA) | $EC_{50}$ (nM) |
| **DA** | 100.00 ± 4.35 | 2426.62 ± 619.46 | 100.00 ± 2.48 | 345.94 ± 60.75 |
| **SKF 81297** | 44.48 ± 1.92**** | 27.48 ± 11.38 | 82.43 ± 2.43** | 9.18 ± 2.50 |
| **SKF 38393** | 21.95 ± 1.68**** | 1180.32 ± 609.05 | 28.32 ± 1.12**** | 121.90 ± 37.36 |
| **DHX** | 109.7 ± 5.78 | 341.98 ± 104.81 | 120.00 ± 4.47*** | 68.23 ± 16.81 |
| **m-DHX** | 70.52 ± 3.44**** | 916.22 ± 285.49 | 120.30 ± 4.53*** | 683.91 ± 166.97 |
| **DOX** | 98.94 ± 5.36 | 572.80 ± 220.80 | 114.40 ± 3.975* | 86.7 ± 23.56 |
| **TAV** | 27.03 ± 2.02**** | 332.66 ± 210.59 | 50.78 ± 1.58**** | 105.44 ± 24.84 |
| **PF-8294** | 53.64 ± 3.60**** | 3698.28 ± 1452.82 | 82.80 ± 2.95** | 2710.19 ± 548.55** |
| **PF-6142** | 39.36 ± 2.97**** | 225.94 ± 149.12 | 76.14 ± 2.80**** | 170.22 ± 47.91 |

Data were presented as mean ± SEM. **, **, ***, ****p<0.05, 0.01, 0.001, 0.0001,* respectively, compared to DA using one-way ANOVA followed by post-hoc Dunnett test.
